# Territoriality and resource availability modulate the effect of conspecific encounters on the foraging behaviours of a mammalian predator

**DOI:** 10.1101/2024.08.14.607927

**Authors:** Jeanne Clermont, Frédéric Dulude-de Broin, Marie-Pier Poulin, Dominique Berteaux

**Affiliations:** Université de Sherbrooke, Sherbrooke (Québec), Canada; Canada Research Chair on Northern Biodiversity, Université du Québec à Rimouski, Rimouski (Québec), Canada; Université Laval, Québec (Québec), Canada; University of Wyoming, Laramie (Wyoming), USA; Center for Northern Studies and Quebec Center for Biodiversity Science

**Keywords:** foraging, home range, intraspecific interactions, movement, territoriality, *Vulpes lagopus*

## Abstract

The probability of encountering conspecifics shapes multiple dimensions of animal behaviour. For example, territorial individuals increase vigilance, scent marking and alarm calling when approaching home range boundaries. Whether territorial predators modify their foraging behaviours with respect to the probability of encountering neighbouring territory owners is poorly understood. However, this could strongly influence the landscape of predation risk and therefore modulate predator-prey interactions. We studied the movements and behaviours of 23 resident Arctic foxes occupying neighbouring home ranges during two years of contrasting resource availability (abundant resources in 2019, scarce resources in 2022) on Bylot Island, Nunavut, Canada. First, based on simultaneous GPS tracking of individuals, we established which individuals used a territory (an exclusive area) by estimating the spatial distribution of the probability of encountering a neighbour within their home range. Second, using GPS and accelerometry data, we evaluated if the probability of encountering a neighbour influenced the spatial distribution of foraging behaviours, and whether this relationship differed between territorial and non-territorial individuals. When resources were abundant, only breeding individuals excluded other foxes from a part of their home range and were thus territorial. When resources were rare, none of the foxes reproduced and all but one were territorial. Non-territorial individuals in 2019 were less likely to cache prey in areas with a high probability of encounter, possibly reducing the risk of cache pilfering. We found no effect of the probability of encounter on the behaviours of the non-territorial individual in 2022. The probability to cache prey of territorial individuals however did not depend on the probability of encountering a neighbour, suggesting they have no benefit of modulating the distribution of their caches according to potential encounters in a partly exclusive area. Our results thus suggest that Arctic foxes use different tactics to secure resources based on their degree of territoriality and the availability of resources. We highlight how the presence of resident, but non-territorial predators, whose home ranges overlap those of their territorial neighbours, may influence the distribution of predation risk by creating zones where predator density is high, potentially influencing predator-prey interactions.

## Introduction

The ability of animals to manage encounters with other individuals is key to all social and interspecific interactions, thus directly influencing reproduction, survival, and ultimately fitness (Silk et al., 2003; Cresswell, 2008; Wey et al., 2008). Consequently, many animal behaviours have evolved to modulate the probability of encountering other individuals. One well known example is how predators and prey modify their habitat use to either find or avoid each other (Sih, 1984; Smith et al., 2019a). Similarly, individuals may avoid conspecifics to decrease resource competition. They for example secure limited resources by becoming territorial, given the benefits of territoriality outweigh the costs (Maher & Lott, 2000; Ord, 2021). A territory represents the area of a home range that is successfully defended by one or several individuals, resulting in the territory owners having exclusive access to the defended area (Brown & Orians, 1970; Powell, 2000; Hinsch & Komdeur, 2017). Territorial individuals will often signal space ownership through visual, auditory or olfactory signals and aggress intruders when encountered (Sillero-Zubiri & Macdonald, 1998; Giuggioli et al., 2011; Tórrez-Herrera et al., 2020).

The intensity of territoriality is often assessed by measuring the amount of overlapping area between neighbouring home ranges (Persson et al., 2010), or the frequency with which individuals adopt territorial behaviours such as scent marking (Fawcett et al., 2013). Furthermore, the degree to which individuals of a given species are territorial can vary among and within populations (Macdonald et al., 1999; McLoughlin et al., 2000; Eide et al., 2004). Ecological variables such as the abundance, predictability and distribution of food resources influence the costs and benefits of territoriality, and therefore partly explain the degree of territoriality (Maher & Lott, 2000; Sells & Mitchell, 2020). Notably, a cost-benefit model developed by Maher and Lott (2000) suggests that individuals should be territorial when food abundance and predictability are intermediate, but non-territorial when food abundance and predictability are below or above some thresholds, which was supported by McLoughlin et al. (2000) who studied variation in home range overlap across brown bear (*Ursus arctos*) populations. Costs and benefits of territoriality may also vary according to predator and competitor densities (Maher & Lott, 2000; Webber et al., 2023), individual characteristics such as sex (Rosell & Thomsen, 2006; Fawcett et al., 2013), or other variables such as the degree of relatedness among neighbours (Persson et al., 2010; Humphries et al., 2021).

The probability of encountering neighbours varies spatially in the home range of territorial animals, being higher near boundaries. Where the probability of encountering a competitor is high, individuals should modify their behaviour to avoid potentially costly physical encounters while still signaling territory ownership more intensively (Schlägel et al., 2017). For example, when located near boundaries, white-faced capuchins (*Cebus capucinus*), which defend group territories, tend to socialise less with each other (Tórrez-Herrera et al., 2020) and travel at lower speed (Noonan et al., 2021), suggesting more vigilance. Also, Ethiopian wolves (*Canis simensis*) and grey wolves (*Canis lupus*) increase scent-marking near boundaries (Sillero-Zubiri & Macdonald, 1998; Zub et al., 2003), while red foxes (*Vulpes vulpes*) spend more time patrolling where artificial scent marks are deposited (Arnold et al., 2011).

An increase in time spent vigilant or in signaling territory ownership when the probability of encountering a competitor is high could further result in fewer time spent foraging (Laundré et al., 2001). Furthermore, probability of encounter may directly affect specific foraging behaviour. For example, in food caching animals that are territorial, such as many canids, felids and mustelids, it may be beneficial to cache food away from competitors to reduce cache pilfering (van der Veen et al., 2020). Better understanding how intraspecific interactions and territoriality shape foraging behaviours could be particularly important for predators, as they generate predation risk landscapes that influence the dynamics of prey populations and community structure (Lima, 2002; Gaynor et al., 2019; Clermont et al., 2021a). For example, in systems where the predator is highly territorial and avoids encountering conspecifics, buffer zones between territories may serve as refuge for prey (Mech, 1977; Lewis & Murray, 1993). In support of this, Anderson et al. (2005) found that reintroduced elks (*Cervus elaphus*) established their home ranges in the periphery of wolf territories and selected areas within their home ranges that were further from wolf territory centers. Prey may thus benefit from fewer predator encounter in places where territorial interference is high and could be further relieved if predators also spend less time foraging in these areas.

Modern tracking techniques help to study predator behaviours as they can locate animals at high frequencies while producing detailed behavioural classification (Nathan et al., 2012; Wilmers et al., 2015). For example, activity level can be identified using GPS data (Patterson et al., 2017), while scent marking (Bidder et al., 2020), killing of prey (Studd et al., 2021) and food caching (Clermont et al., 2021b) may be identified using accelerometry. Estimating where encounters between individuals occur is however challenging, limiting our ability to assess how probabilities of encounter affect behaviours (Noonan et al., 2021). So far, most research has compared how animals behave in and out of overlapping areas or with respect to the distance to territory borders (assuming these variables are good surrogates for encounter probability), instead of directly estimating the spatial distribution of encounter probabilities (Rosell & Thomsen, 2006; Tórrez-Herrera et al., 2020). To address this methodological gap, Noonan et al. (2021) proposed a statistical framework using tracking data and home range estimation to evaluate the spatial distribution of the probability of encounter, termed the conditional distribution of encounters (CDE). The CDE quantifies where in space encounters are likely to occur within an individual’s home range. Combining behavioural classifications to the CDE should allow to better understand how interactions with neighbours affect behaviours of territorial animals, further improving our understanding of important ecological processes.

We used GPS tracking and accelerometry to monitor the movements and behaviours of Arctic foxes living in a large Greater snow goose (*Anser caerulescens atlanticus*) colony on Bylot Island (Nunavut, Canada) (Clermont et al., 2021a, 2021b) during two years of contrasting resource availability. At this site, foxes are socially monogamous, offer biparental care, and most home ranges are occupied by a mated pair (Cameron et al., 2011). Food caching is an important dimension of the foraging ecology of Arctic foxes, which cache the majority of the goose eggs they collect (Careau et al., 2007). Our first objective was to quantify the degree of summer territoriality of each studied individual by mapping the probability of encountering conspecifics other than the mate within the home range, and to evaluate whether fox territoriality depends on the availability of food resources. Following Maher and Lott’s (2000) model predictions, we predicted that fewer foxes would be territorial when resources are scarce (P1). Our second objective was to test the hypothesis that the probability of encountering a neighbour influences the spatial distribution of foraging behaviours within home ranges, and test whether territoriality modulates this relationship. As Arctic foxes may attack other foxes foraging within their territory and perform cache pilfering (Samelius & Alisauskas, 2000; Careau et al., 2007), we predicted (P2) that foxes should be less likely to hunt and cache prey where the probability of encountering a neighbour and the risk of cache pilfering is high. Lastly, we predicted (P3) that the probability of encountering a neighbour should more strongly affect the foraging and caching behaviour of non-territorial than territorial foxes, since the latter individuals might focus on excluding other foxes from their territory rather than avoiding encounters.

## Materials and methods

### Study system

We worked in 2019 and 2022 in the southwest plain of Bylot Island (72°53′ N, 79°54′ W) in Sirmilik National Park of Canada (Nunavut), where the Arctic fox is the main terrestrial predator. Arctic foxes at this site are highly range resident (i.e., they use a stable home range), they usually share a home range with their mating partner and most individuals show low home range overlap with their neighbours (Grenier-Potvin et al., 2021). They bark and scent mark to indicate territory ownership, and they chase intruders (Eberhardt et al., 1982). In addition to selecting habitats suitable to their main prey, they avoid home range borders, potentially to minimise interactions with their neighbours (Grenier-Potvin et al., 2021).

On Bylot, Arctic foxes rely mostly on small prey, such as lemmings (*Lemmus trimucronatus* and *Dicrostonyx groenlandicus*). They also feed on greater snow goose eggs when their home range overlaps Bylot’s large colony of more than 20,000 nesting pairs (Bêty et al., 2001). In 2022 however, nesting goose density was unusually low throughout the whole colony (density estimated from random plots: 128 nests/km^2^ in 2022, 438 nests/km^2^ in 2019 and an average of 248 nests/km^2^ between 2000-2019; Cadieux et al., 2023). Foxes also opportunistically prey on the nests of other ground nesting birds (Duchesne et al., 2021). Lemming density fluctuates cyclically (Gruyer et al., 2008), and was moderate in 2019 and very low in 2022 as determined by capture-recapture methods (Fauteux et al., 2015; Duchesne et al., 2021; Cadieux et al., 2023). The snow goose incubation period lasts 23 days from mid-June to early July, during which foxes collect eggs for later consumption (Samelius et al., 2007). In years of low to moderate lemming densities, goose eggs represent the majority of prey collected by foxes during the goose incubation period (Careau et al., 2007). Foxes cache up to 90% of the goose eggs they collect, and caches are located 85 m (median) from the nest (Careau et al., 2007). They also recache ca. 60% of goose eggs, and recovery rate (for recaching or consumption) is highest at the end and after the goose incubation period (Careau et al., 2008).

Furthermore, access to the snow goose colony influences fox probability of reproduction (Chevallier et al., 2020). Despite interindividual variations, the phenology of fox reproduction largely coincides with that of geese, with cubs remaining in the den during the goose egg incubation period, and progressively emerging and becoming independent from the den during goose brooding (Grenier-Potvin et al., 2021). Due to low densities of both lemmings and goose nests in 2022, none of the studied foxes reproduced that year, contrary to 2019 when many foxes reproduced.

### Arctic fox tracking

In May and June, we captured 13 foxes in 2019 and 10 foxes in 2022 in the snow goose colony using Softcatch #1 padded leghold traps (Oneida Victor Inc. Ltd., Cleveland, OH, USA). We determined the individual’s sex from genital characteristics and fitted them with four coloured-ear tags and a GPS-accelerometer collar (95 g, ca. 4% of body mass; Radio Tag-14, Milsar, Romania) for subsequent identification and tracking. Arctic fox capture techniques and immobilization procedures were approved by the UQAR Animal Care Committee (CPA-64-16-169 R3) and field research was approved by the Joint Park Management Committee of Sirmilik National Park of Canada (SIR-2018-28021).

In 2019, the 13 tracked foxes consisted of six neighbouring pairs plus one individual, and their home ranges formed our 2019 study area (Figures 1A and 1B). In 2022, the 10 tracked foxes included one pair for which both individuals were tracked, and eight pairs for which only one individual of the pair was tracked, and their home ranges formed our 2022 study area (Figures 1C and 1D). We determined reproductive status of individuals based on whether automated cameras recorded cubs at individuals’ den (Cameron et al., 2011). Daily observations and automated cameras at fox dens confirmed that all foxes within our study area were tracked in 2019 and at least one individual per pair in 2022. We collected one GPS location every 4 minutes and one 30-sec burst of accelerometry (50 Hz) every 4.5 minutes in 2019 and every 4 minutes in 2022 (Clermont et al., 2021b). We divided the 30-sec bursts of accelerometry into ten 3-sec sequences and, using behavioural observations and a random forest algorithm, we assigned to each sequence one of four behaviours: running, walking, digging and motionless (see analytical details in Clermont et al., 2021b). During the goose incubation period, digging behaviour of foxes is mainly associated with goose egg caching, but may also occasionally reflect caching and handling of other prey species as well as cache recoveries (Clermont et al., 2021b). As most captured prey are cached (Careau et al., 2007), fox digging behaviour is more generally a proxy of foraging activity.

**Figure 1.**
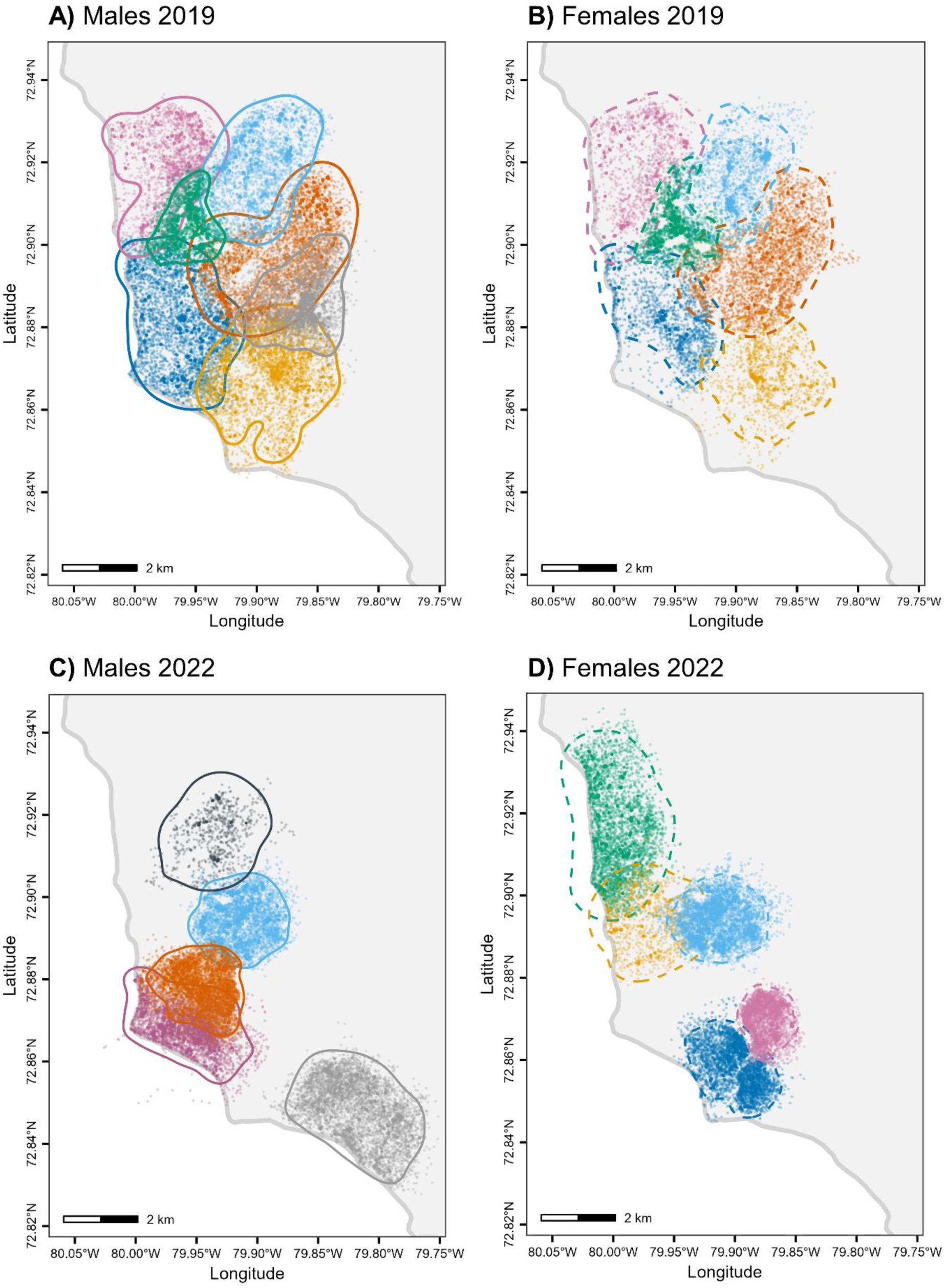
GPS locations (dots) of male and female Arctic foxes tracked from 11 June to 4 July 2019 when resources were abundant (n = 67,109, panels A and B) and from 19 June to 12 July 2022 when resources were scarce (n = 47,631, panels C and D) at a 4-min fix interval on Bylot Island, Nunavut, Canada. Solid (males) and dashed (females) lines are 95% home range contours obtained through auto-correlated kernel density estimation. For each year, the same colours are used for members of a given fox pair. Home ranges are located near the coast of Bylot island shown by the thick grey line.

### Data analysis

#### 1) Estimation of homes ranges

To define Arctic fox summer home ranges, we used GPS data collected from 11 June to 4 July 2019, and 19 June to 12 July 2022, corresponding to goose egg incubation periods. Prior to home range estimation, we confirmed home range residency for each individual using variogram analysis (Fleming et al., 2014), and excluded location data resulting from extra-territorial trips as it affects home range estimation (Calabrese et al., 2016). Indeed, during the study period, some individuals made a few extra-territorial trips going up to 25 km away from the center of the home range. These “outliers” were identified by evaluating distances between points and home range core, and a cut-off distance was determined visually for each individual (see Appendix A for further details). We excluded 11%, 9%, 6% and 4% of datapoints for four individuals that performed extra-territorial trips (three individuals in 2019 and one in 2022). The remaining individuals showed high range residency and we excluded < 1% of their GPS locations.

We then fitted range-resident continuous-time movement models to the data of each individual (model selection resulted in the Ornstein-Uhlenbeck Foraging (OUF) process used for each individual), to control for autocorrelation in both speed and location (Fleming et al., 2014; Calabrese et al., 2016). Then, we estimated home range areas using auto-correlated kernel density estimation (AKDE; Fleming et al., 2015). AKDE is more accurate than other home range estimators for autocorrelated location data (Noonan et al., 2019). The complete workflow we used is detailed in Silva et al. (2021), and was performed using the package ctmm (v0.6.2, Calabrese et al., 2016) in R 4.1.0 (R Development Team, 2021).

#### 2) Estimation of probability of encounter among neighbours

Following Noonan et al. (2021), we estimated the conditional distribution of encounters (CDE) for each year separately (package ctmm v0.6.2). The CDE estimates the spatial distribution of encounter events in the environment as the normalized product of AKDE home range estimates.

The probability of encounter estimated by the CDE thus represents the probability that more than one individual uses a specific area, but not necessarily at the same time. A CDE value of 1 at a given location indicates there is a 100% chance that more than one individual use that location, and thus that the probability of encounter is high, and a value of 0 that only one individual uses that location, meaning individuals should never encounter each other at that location. Since we were interested in encounters among neighbours, we excluded interactions between mates within their shared home range. We then determined the degree of territoriality of each fox and compared the proportion of territorial individuals between years (P1). A fox was considered territorial if at least some part of its home range was outside of the 95% CDE (Noonan et al., 2021), excluding areas within 500 m of boundaries of study areas. Indeed, some interactions inevitably occurred at home range boundaries located at the edge of the 2019 and 2022 study areas (but not near the coastline, Figures 1 and 2), where neighbouring foxes were not tracked, leading to a local underestimation of CDE values. Using a 1 km buffer instead of a 500 m one would have led to the same classification (territorial yes/no) of all individuals but one in 2022 (female BJOV; see Appendix B).

**Figure 2.**
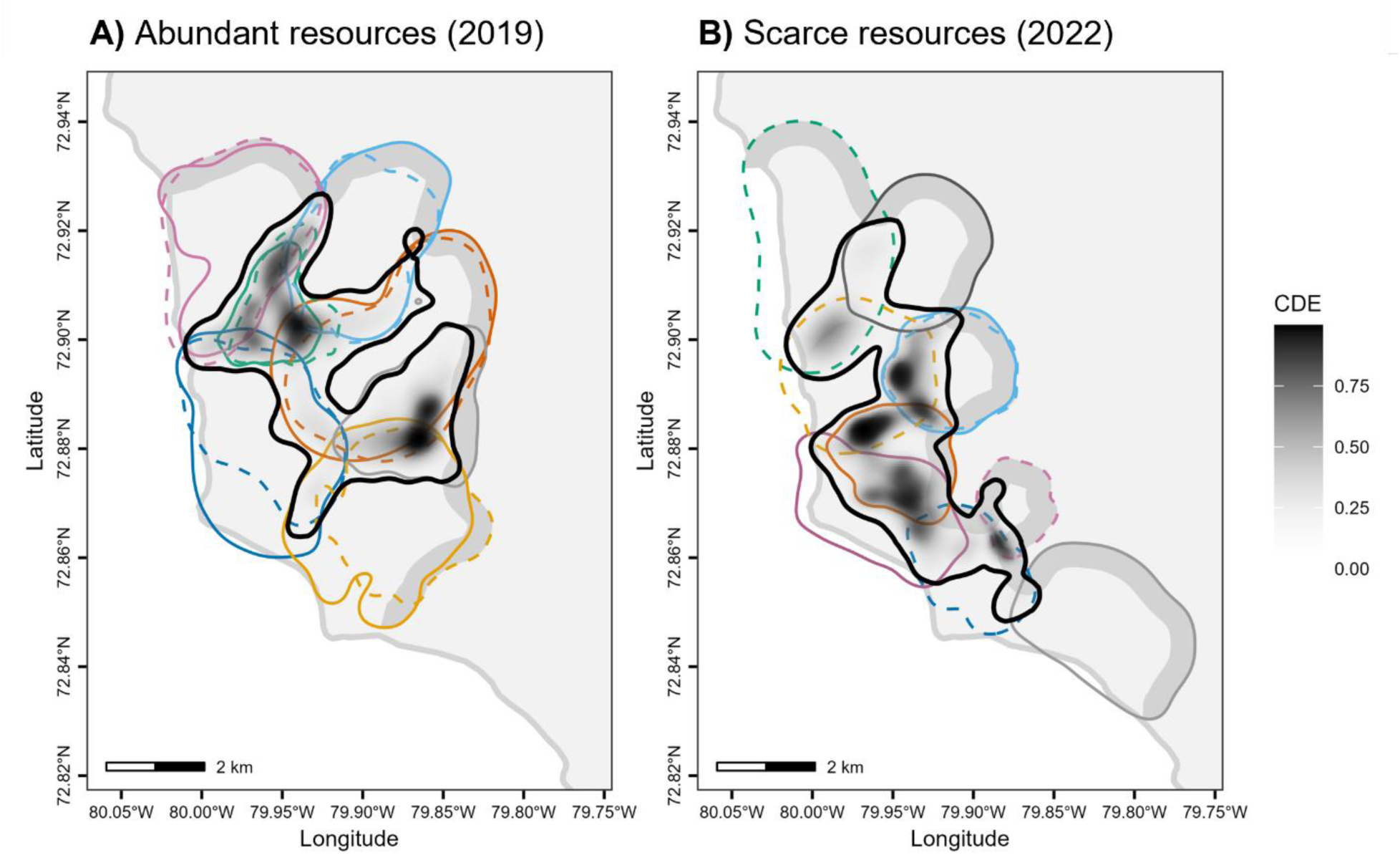
Conditional distribution of encounters (CDE) among A) 13 Arctic fox neighbours in 2019 when resources were abundant and B) ten Arctic fox neighbours in 2022 when resources were scarce, tracked with GPS, excluding interactions between pair members. The thick black line is the 95% CDE contour and coloured lines are 95% AKDE contours. The grey shaded area represents the 500 m inner buffer of study area boundaries where interactions with other foxes inevitably occur. Males and females are represented by solid and dashed lines, respectively, and members of a given pair are identified by the same colour within each year.

#### 3) Arctic fox prey caching behaviour

We used our accelerometry behavioural classification to determine whether individuals engaged in digging (indicating prey caching, cache recovery events and more generally foraging activity) during each 30-sec accelerometry burst. At least one out of the ten 3-sec accelerometry sequences had to be assigned to the digging behaviour to consider that the fox engaged in digging in that burst. Each 30-sec burst was then associated to the closest GPS location, provided the time stamp of the GPS location was within 30 sec of the start or end of the burst. Bursts occurring at less than 50 m of a den were excluded as digging may then reflect den maintenance rather than prey caching.

We excluded from analyses all accelerometry bursts occurring within 500 m of the boundaries of study areas, since interactions with unknown foxes may then occur (Appendix B). To verify whether buffer size influences results, we 1) repeated the analyses described in the next section by excluding the individual of 2022 that would have been considered as non-territorial when using a 1 km buffer (female BJOV), and 2) repeated our analyses by excluding locations falling within a 1 km buffer instead of a 500 m one (Appendix B). In both cases, we obtained similar results.

#### 4) Effect of the probability of encountering a neighbour on fox behaviours

To test the influence of the likelihood of encountering a neighbour on the probability of engaging in foraging behaviours (objective 2), we used generalized additive mixed models (GAMM) with a binomial error distribution and logit link function (“gam” function of the R package mgcv, v1.9-0, Wood, 2017) and modelled fox probability of engaging in digging (0 = no digging, 1= digging; P2) with respect to the probability of encounter. As not all foxes experienced the same range of probability of encounter, we centered the probability of encounter values to a mean of 0 within each fox home range. We modeled 2019 and 2022 data separately. In both models, we included as fixed effect a categorical variable representing whether the fox was territorial (yes/no), which we included in interaction with the probability of encounter, to test whether the degree of territoriality modulated the effect of the probability of encounter on the probability of caching (P3). In 2019, we also controlled for nesting goose density (individual geese/ha), a proxy for goose nest density estimated from detailed field surveys (Grenier-Potvin et al., 2021), as nesting goose density was previously shown to affect fox probability to dig (Clermont et al., 2021b). We did not perform detailed field surveys in 2022 as nest density was very low throughout the study area, therefore we could not include nesting goose density as a covariate in the 2022 models. We included individual sex as another fixed effect, as well as fox identity and Julian date as random intercepts. To address temporal autocorrelation in our datasets, we also added to all models a cyclic cubic spline function of the numeric time of the day and a thin plate regression spline function of the Julian date, with 10 basis dimensions for each function (Pedersen et al., 2019). All continuous covariates were centered and standardised to facilitate the interpretation of model estimates (Schielzeth, 2010). We verified model assumptions and independence of residuals using the R package DHARMa (v0.4.3, Hartig, 2021), and observed significant deviations in models’ residuals. We solved these issues by using a subset of data that included 150 randomly chosen observations by individual for both models, and by keeping all covariates in the models even when effects were not significant. Results are expressed using evidence-based language (Muff et al., 2021) where p-value ≥ 0.1 suggests no evidence, p-value =]0.05, 0.1] suggests weak evidence, p-value =]0.01, 0.05] suggests moderate evidence, p-value =]0.01, 0.001] suggests strong evidence and p-value < 0.001 suggests very strong evidence.

## Results

### Conditional distribution of encounters in fox home ranges

Home range areas averaged 7.93 ± 3.15 (SD) km^2^ (Figure 1, n = 23). We found no evidence that home range areas differed between sexes (t-test: t = 0.53, p = 0.600, average male home range area = 8.27 ± 3.12 (SD) km^2^, n = 12, average female home range area = 7.56 ± 3.28 (SD) km^2^, n = 11). However, there was weak evidence that home ranges were smaller in 2022 than in 2019 (t-test: t = 2.07, p = 0.051, average home range area in 2019 = 9.03 ± 3.12 (SD) km^2^, n = 13, average home range area in 2022 = 6.51 ± 2.70 (SD) km^2^, n = 10).

As expected, higher CDE values occurred where home ranges of tracked neighbours overlapped (Figure 2). When resources were abundant (2019), three individuals (green female RMJJ, green male BORR and grey male BBJO) were non-territorial as they had all of their home range located within the 95% CDE contour if we ignore 500 m study area edges (Figures 2A and 3A). These three foxes did not reproduce in 2019. In contrast, all other ten individuals were territorial, having a considerable area of their home range outside the 95% CDE contour, where the probability of encounter with a neighbour is null (Figures 2A and 3A). These ten individuals reproduced in 2019. When resources were rare (2022), all individuals but one (dark orange male BOBB) were territorial (Figures 2B and 3B). None of the foxes reproduced in 2022. We thus find no support for an effect of resource abundance on territoriality (P1) as 10/13 and 9/10 individuals were territorial in 2019 (high resource abundance) and 2022 (low resource abundance), respectively.

**Figure 3.**
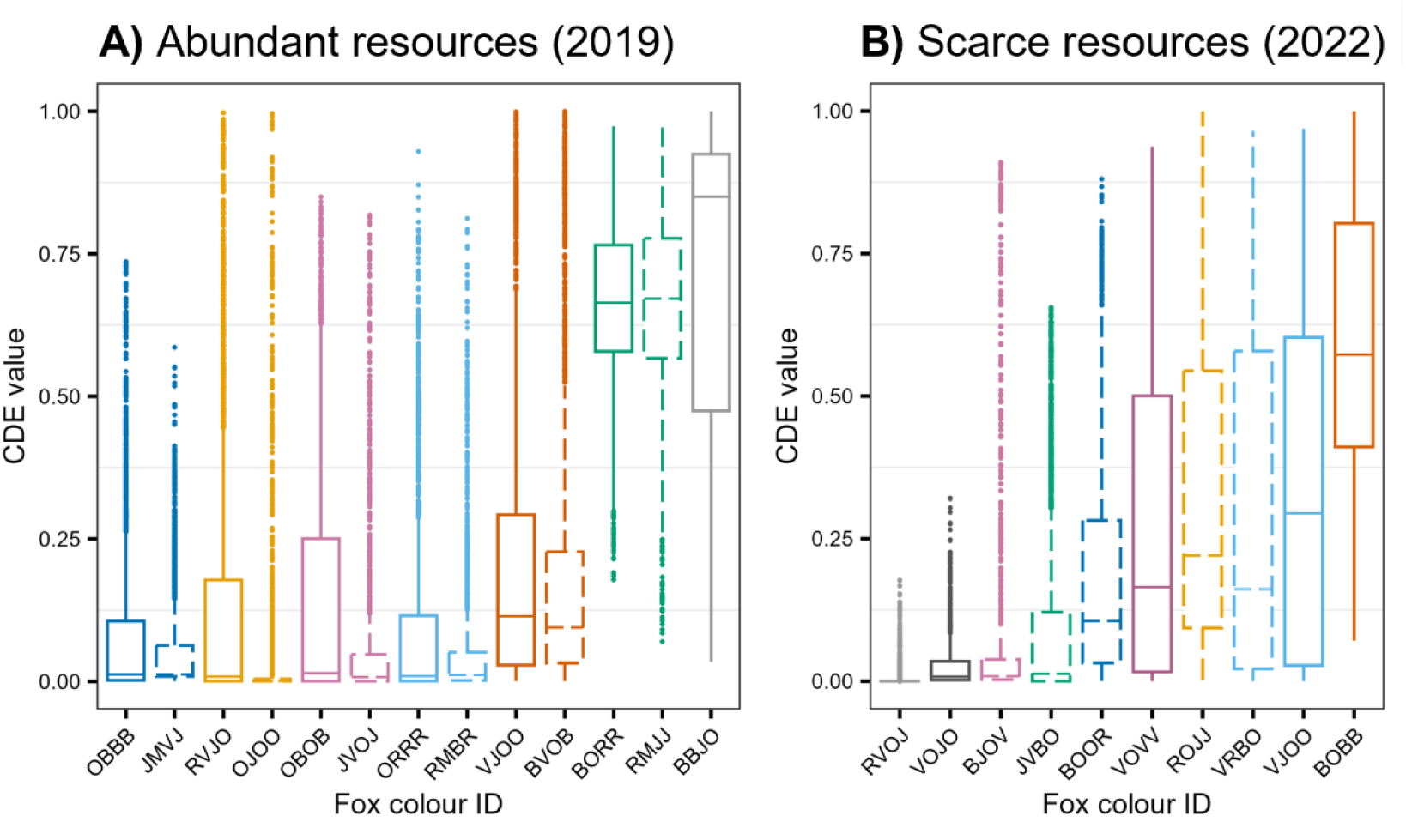
CDE values (probability of encountering a neighbour) observed at each GPS location from Figure 1, for each fox in A) 2019 and B) 2022. Fox colour IDs reflect the unique combinations of colours of the four ear tags (e.g., OBBB: orange-blue-blue-blue). Boxplot colours used for each individual are the same as in Figures 1 and 2. Males and females are represented by solid and dashed lines, respectively, and each year’s members of a given pair are identified by the same colour. Boxplots show first quartile, median, and third quartile. Lower and upper whiskers extend, respectively, to the lowest and highest values within the interquartile range multiplied by 1.5. Points represent values outside this range.

### Effect of the probability of encountering a neighbour on fox caching behaviour

When resources were abundant (2019), non-territorial foxes were more likely to engage in digging (strong evidence) in places with low encounter probability (Table 1; Figure 4A). However, when resource were scarce (2022), we found no evidence that the probability of encounter influenced digging behaviour of the only non-territorial individual (Table 1; Figure 4B). As for territorial foxes, we did not find evidence that digging behaviour was influenced by the probability of encountering a neighbour in either year (Table 1; Figure 4). Thus, when resources were abundant, the probability of encountering a neighbour only affected the behaviours of non-territorial individuals, and all individuals were unaffected when resources were scarce (Figure 4), providing partial support for an effect of encounter probability on foraging behaviours (P2) and for a larger influence of encounter probability in non-territorial foxes (P3). We also found very strong evidence that nesting goose density has a positive effect on the propensity to dig in the year where that variable could be included (2019; Table 1). There was no evidence of an effect of sex in both years (Table 1). To ensure results were robust despite the low sample size, we ran our models on different subsets of the dataset and observed slight changes in coefficient values, but no differences in the direction and significance of effects.

**Figure 4.**
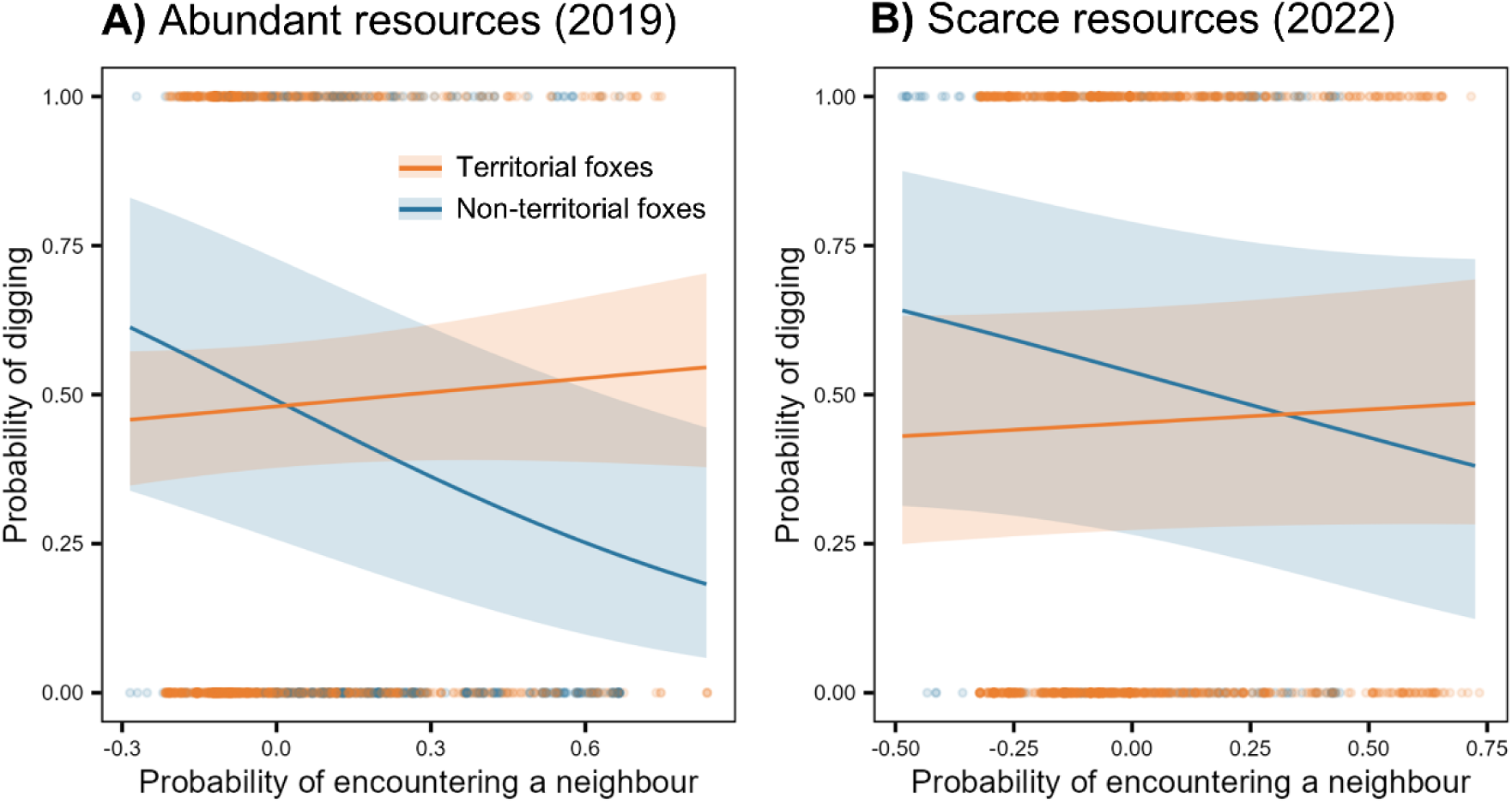
Predicted effects of the probability of encountering a neighbour on probability of fox digging (0 = no digging, 1= digging) in A) 2019 when resources were abundant (n = 1950 accelerometry bursts) and B) 2022 when resources were scarce (n = 1500 accelerometry bursts). The probability of encountering a neighbour was obtained from conditional distributions of encounters (Figure 2). Orange dots and lines refer to territorial individuals (n = 10 foxes in 2019 and 9 foxes in 2022), and blue dots and lines refer to non-territorial individuals (n = 3 foxes in 2019 and 1 fox in 2022). Shaded areas represent 95% confidence intervals.

**Table 1.**
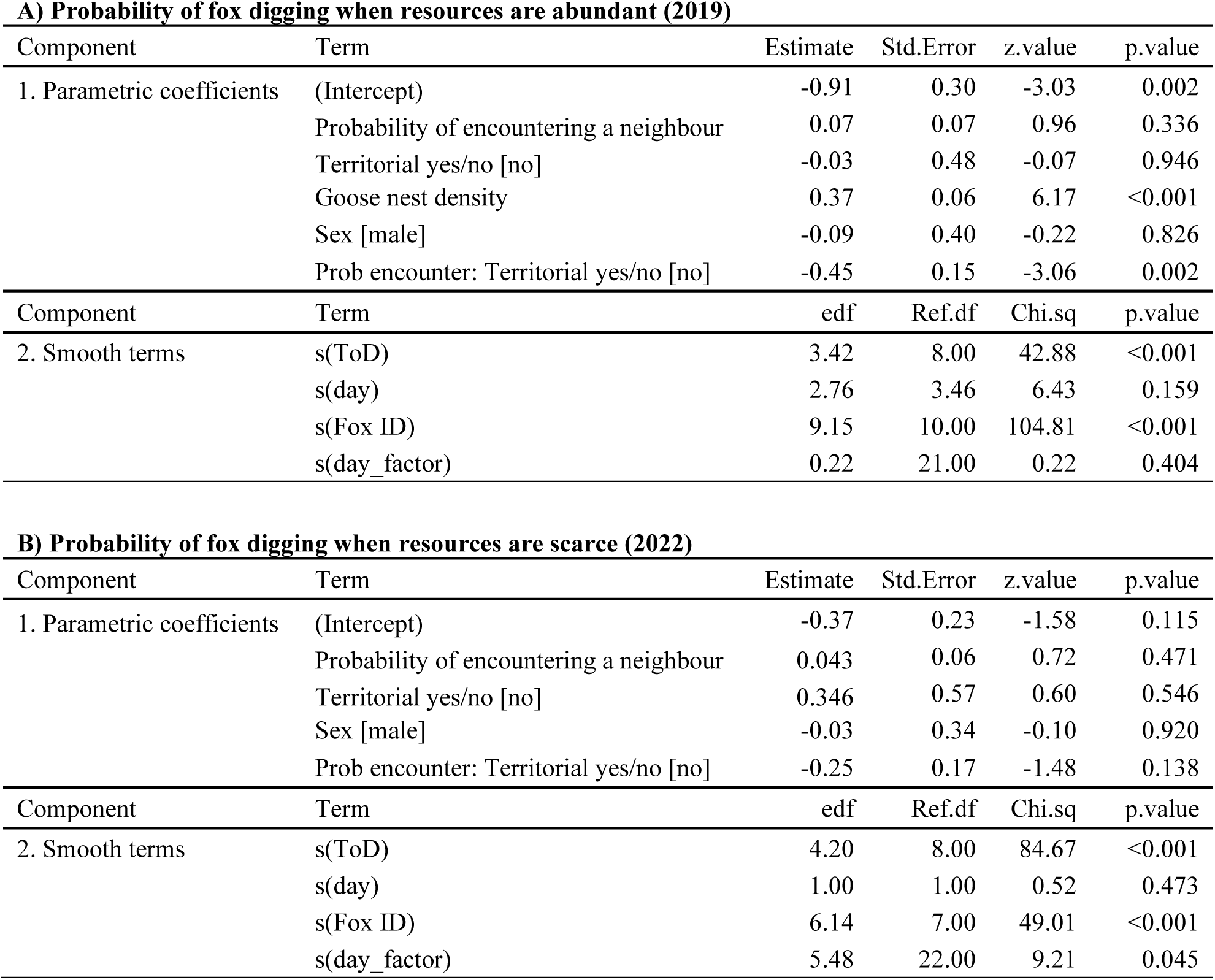
Estimates of generalized additive mixed models predicting the probability of fox digging in A) 2019 when resources were abundant (n = 1950 accelerometry bursts) and B) 2022 when resources were scarce (n = 1500 accelerometry bursts). Models had a binomial error distribution with a logit-link function. We included as parametric terms (fixed effects) the probability of encountering a neighbour, whether the fox was territorial, an interaction between the probability of encountering a neighbour and whether the fox was territorial, the individual’s sex and goose nest density (2019 only). We included splines for numeric time of day (ToD) and Julian date (day), and the random effects fox identity (Fox ID) and Julian date (day_factor).

| **A) Probability of fox digging when resources are abundant (2019)** |  |  |  |  |  |
| --- | --- | --- | --- | --- | --- |
| Component | Term | Estimate | Std.Error | z.value | p.value |
| 1. Parametric coefficients | (Intercept) | -0.91 | 0.30 | -3.03 | 0.002 |
|  | Probability of encountering a neighbour | 0.07 | 0.07 | 0.96 | 0.336 |
|  | Territorial yes/no [no] | -0.03 | 0.48 | -0.07 | 0.946 |
|  | Goose nest density | 0.37 | 0.06 | 6.17 | <0.001 |
|  | Sex [male] | -0.09 | 0.40 | -0.22 | 0.826 |
|  | Prob encounter: Territorial yes/no [no] | -0.45 | 0.15 | -3.06 | 0.002 |
| Component | Term | edf | Ref.df | Chi.sq | p.value |
| 2. Smooth terms | s(ToD) | 3.42 | 8.00 | 42.88 | <0.001 |
|  | s(day) | 2.76 | 3.46 | 6.43 | 0.159 |
|  | s(Fox ID) | 9.15 | 10.00 | 104.81 | <0.001 |
|  | s(day_factor) | 0.22 | 21.00 | 0.22 | 0.404 |

| **B) Probability of fox digging when resources are scarce (2022)** |  |  |  |  |  |
| --- | --- | --- | --- | --- | --- |
| Component | Term | Estimate | Std.Error | z.value | p.value |
| 1. Parametric coefficients | (Intercept) | -0.37 | 0.23 | -1.58 | 0.115 |
|  | Probability of encountering a neighbour | 0.043 | 0.06 | 0.72 | 0.471 |
|  | Territorial yes/no [no] | 0.346 | 0.57 | 0.60 | 0.546 |
|  | Sex [male] | -0.03 | 0.34 | -0.10 | 0.920 |
|  | Prob encounter: Territorial yes/no [no] | -0.25 | 0.17 | -1.48 | 0.138 |
| Component | Term | edf | Ref.df | Chi.sq | p.value |
| 2. Smooth terms | s(ToD) | 4.20 | 8.00 | 84.67 | <0.001 |
|  | s(day) | 1.00 | 1.00 | 0.52 | 0.473 |
|  | s(Fox ID) | 6.14 | 7.00 | 49.01 | <0.001 |
|  | s(day_factor) | 5.48 | 22.00 | 9.21 | 0.045 |

## Discussion

Using high throughput tracking technologies, we estimated the spatial distribution of the probability of encounter among Arctic foxes using neighbouring home ranges, during two years of contrasted resource abundance varying from high (2019) to low (2022). We benefited from a natural situation where most, but not all individuals were territorial, thus enabling us to evaluate the effect of territoriality on the spatial distribution of foraging behaviour. When resources were abundant, non-territorial individuals responded to the variation in the probability of encountering a neighbour by shifting the distribution of prey caching away from the places where encounters were most likely to occur. The only non-territorial individual monitored when resource abundance was low however did not respond to variation in the probability of encounter in its home range. The probability of encountering a neighbour also did not influence territorial individuals’ prey caching behaviour. Despite modern technology, it remains highly challenging to simultaneously track the movements and foraging behaviours of many neighbouring vertebrate predators at high resolution, in a natural setting and over several weeks. Our sample of 13 individuals in 2019 and 10 individuals in 2022, each tracked for more than three weeks, is therefore of high value. Our study provides valuable insights into how intraspecific interactions may influence the spatial distribution of predator foraging.

### The degree of territoriality varies among foxes

Variation in the distribution, abundance and predictability of resources often explains among-individual differences in territoriality (Maher & Lott, 2000; Sells & Mitchell, 2020). For example, Eide et al. (2004) observed that in Arctic foxes of Svalbard, high home range overlap occurred near the coast where seabird colonies are concentrated. On the other hand, little home range overlap occurred inland where reindeer (*Rangifer tarandus*) carcasses are scattered and unpredictable. In our study, we do not suspect that variation in prey distribution, abundance or predictability explains among-individual differences in territoriality within years, as all studied foxes lived near the center of the goose colony where nest abundance is usually high and habitats favourable to lemmings. However, we used two contrasted years of data, with one year of moderate lemming abundance and typical goose density for this study area (2019), and one year of low lemming abundance and low goose density (2022). In 2019, although resource abundance was high, not all foxes reproduced and used a territory. This suggests that using a territory may be necessary to secure enough resources to ensure the survival of young, even when food is abundant (Webb et al., 2012; López-Bao et al., 2014). On the other hand, for non-breeding individuals, the costs of defending a territory may have been too high compared to the benefits when resources are abundant, as predicted by Maher and Lott’s (2000) cost-benefit model of territoriality.

Although resources were scarce in 2022, all foxes but one could exclude neighbours from part of their home range, which seems to contradict Maher and Lott’s (2000) model, that predicts no territoriality when resources are very rare. This could however indicate that food abundance inside the colony in 2022 was still not low enough for territoriality not to be beneficial. In fact, foxes in the colony might rely on stored eggs from the previous year (Samelius et al., 2007), which could attenuate the effects of low resources. However, although high abundance of resources may permit non-breeding individuals to forego being territorial, defending a territory may remain beneficial under low or very low resource abundance, especially when the abundance of resources varies cyclically through time. Similarly, López-Bao et al. (2014) observed that home range overlap among Iberian lynx *(Lynx pardinus*) remains low across a gradient of prey availability. This obstinate strategy (sensu von Schantz, 1984), i.e., maintaining a territory even when resources are rare, could be beneficial if it allowed to secure a home range in areas where resources are usually high, such as the snow goose colony on Bylot Island. Therefore, it may remain beneficial for foxes to defend a territory in the goose colony even during years of low goose nesting density, to maintain territory ownership and maximise chances of future reproduction (López-Bao et al., 2014).

### Variable effect of the probability of encountering a neighbour on Arctic fox foraging behaviours

Foraging and caching prey in places where the probability of encountering a neighbour is high can increase the risk of injury or cache pilfering (Samelius & Alisauskas, 2000; van der Veen et al., 2020). Therefore, Arctic foxes should be more likely to forage and cache prey where they are less likely to encounter neighbours. However, such behaviour can be influenced by the intrinsic and environmental constraints faced by individuals. We found that resource abundance and the level of territoriality influenced how Arctic foxes responded to variation in the probability of encountering a neighbour. When resources were abundant, non-territorial foxes had a higher probability to dig where the probability of encountering neighbours was low. For these individuals without exclusive home range, foraging and caching prey where the risk of cache pilfering is low may increase foraging efficiency. Although during goose incubation period, digging events should mostly reflect prey caching, we cannot differentiate caching events from prey captures, cache recoveries and recaching events (Clermont et al., 2021b). Therefore, our results may also indicate that non-territorial individuals could favour recaching away from their neighbours, in safer sites. Caching and recaching prey in “out of view” sites are in fact common cache protection strategies in other species such as corvids (Dally et al., 2006). Careau et al. (2007) also showed that Arctic fox recaches goose eggs away from were they were collected, potentially to secure eggs in sites where chances of pilfering are lower (e.g., closer to the den). The non-territorial individual of 2022 however cached prey independently of its probability of encountering neighbours, suggesting that when resources are scarce, non-territorial foxes may not afford to cache prey only in specific areas of the landscape, but this result should be confirmed with a larger sample of individuals.

Territorial foxes did not adjust caching behaviours to the probability of encountering a neighbour in either year of contrasted resource abundance. By defending a territory, these individuals had null encounter probability in substantial parts of their home range. This likely reduced the need to modulate foraging and caching behaviours, a potential benefit of establishing a territory. Still, these individuals could have withheld from foraging and caching in the few parts of their home range where encounter probability was not null. However, our results suggest that they did not, potentially because focusing on maintaining and defending their territory is more important than avoiding risk of injury or cache pilfering by conspecifics. Indeed, territorial individuals are likely more dominant and better suited to win an encounter than non-territorial individuals (Stamps & Krishnan, 2001). It might therefore be beneficial for these individuals to actively use all parts of their home range at the risk of injury and cache pilfering rather than leaving some areas undefended. Our results suggest that territoriality could be enough to secure the necessary resources for survival and reproduction.

In addition to spatial adjustments of behaviours, Arctic foxes may modify their foraging behaviours temporally to avoid encounters, which the CDE did not allow to evaluate (Noonan et al., 2021). Similarly, prey species access resources in high-predation risk areas at times of the day when predators are less active (Kohl et al., 2018; Smith et al., 2019b). Future studies should include temporal partitioning when estimating the probability of encounter to help uncover the different tactics territorial animals use to avoid intraspecific interactions while foraging.

### Implications for predator-prey interactions

Arctic foxes have important top-down effects in the tundra and generate multiple predator-mediated interactions among prey (Bêty et al., 2002; Legagneux et al., 2012; Duchesne et al., 2021). For example, nest survival of American golden plover (*Pluvialis dominica*) is lower within the snow goose colony, where foxes use smaller home ranges and are thus present at higher densities (Dulude-de Broin et al., 2023). At fine spatial scales, Arctic fox movements generate a predation risk landscape influencing the behaviour and nest distribution of several migratory birds (Clermont et al., 2021a). It is therefore of strong interest to identify the factors explaining Arctic fox density and where they choose to forage. In this study, we observed that the degree of territoriality and food abundance influence fox response to intraspecific interactions. When resources are abundant, non-territorial individuals avoid caching prey where the probability of encountering a neighbour is high. As most captured prey are also cached (Careau et al., 2007), our results suggest non-territorial individuals forage less intensively in areas likely used by other individuals. However, there was no effect of the probability of encounter on the caching behaviour of territorial individuals (i.e., most individuals), suggesting intraspecific interactions may not have a strong effect on the spatial distribution of predation risk. Interestingly, although most Arctic foxes were territorial, we observed no unused areas between adjacent home ranges, but instead high probability of encounters in overlapping areas (Figures 1 and 2). As such, the spatial configuration of fox home ranges should not lead to buffer zones of low predation risk between territories as observed in some wolf-ungulate systems (Mech, 1977; Anderson et al., 2005). Conversely, the presence of non-territorial individuals, whose home ranges overlap those of their territorial neighbours, creates zones where fox density is high, which can increase the risk of predation (Clermont et al., 2021a; Dulude-de Broin et al., 2023). As such, we hypothesise that the greatest risk of predation should be located within the home range of non-territorial individuals, while the lowest predation risk should be found in defended territories where the probability of encounter among neighbouring foxes is null.

### Conclusion

We found that not all Arctic foxes on Bylot Island used a territory. We highlight that because costs and benefits of territoriality may differ among individuals in a population, alternative behavioural tactics may emerge from non-territorial individuals to secure resources. Including the conditional distribution of encounters into habitat selection analyses would further help to better understand how different individuals deal with both the physical and social environments when using their habitat, and how they compromise between the needs to acquire valuable resources and avoid competitors. Finally, a spatial configuration in which the home ranges of non-territorial individual predators overlap those of their territorial neighbours may influence the distribution of predation risk, which could modulate the distribution and behaviour of prey and the structure of communities.

## Data Accessibility Statement

Arctic fox GPS and accelerometry data are available through the Movebank Data Repository (Movebank Study ID 1241071371 and 2602389249).

## Author Contributions

All authors conceived the study and collected the data. JC performed the analyses with help from FDB and MPP. JC wrote the first version of the manuscript, which all authors commented and revised. All authors approved the final version.

## Acknowledgements

We thank A. Grenier-Potvin, R. Gravel, G. Roy, L.-P. Ouellet, M. Beaudoin and M. Archambault for field work, M. Noonan for advice on spatial analyses, and the community of Mittimatalik for support. Funding was received from (alphabetical order) Canada Foundation for Innovation, Canada Research Chairs Program, Fonds de Recherche du Québec – Nature et technologies (FRQNT), Natural Sciences and Engineering Research Council of Canada (NSERC), Network of Centers of Excellence of Canada ArcticNet, Northern Scientific Training Program (Polar Knowledge Canada), Parks Canada Agency, Polar Continental Shelf Program (Natural Resources Canada), and Weston Family Foundation.

## Appendix A: Workflow to exclude GPS locations associated to extra-territorial trips for home range estimation

Some individuals performed extra-territorial trips (Figure A1), which affected home range estimation. We considered these GPS locations associated to extra-territorial trips as outliers and identified them by calculating the distance between the core of the home range and each GPS location in the R package ctmm with the function ‘outlie’ (Calabrese et al., 2016). Figure A2 shows distance values for the individual BORR in 2019 (green male on Figure 1A of the main text) and VOVV in 2022 (dark purple male on Figure 1C of the main text). Zooming in on lower distance values helped to determine cut-offs for individuals travelling far from their home range core (Figure A3). For BORR, we removed GPS locations >1700 m from the home range core, which removed 9% of its original dataset. For VOVV, we removed GPS locations >2300 m from the home range core, which removed 11% of its original dataset. These thresholds were determined visually so that extra-territorial trips disappeared from the spatial representation of the individual’s movements (Figure A4). We also removed 4% of GPS locations for RMJJ in 2019 (green female on Figure 1B), 6% for BBJO in 2019 (grey male on Figure 1A), and < 1% for all others.

**Figure A1.**
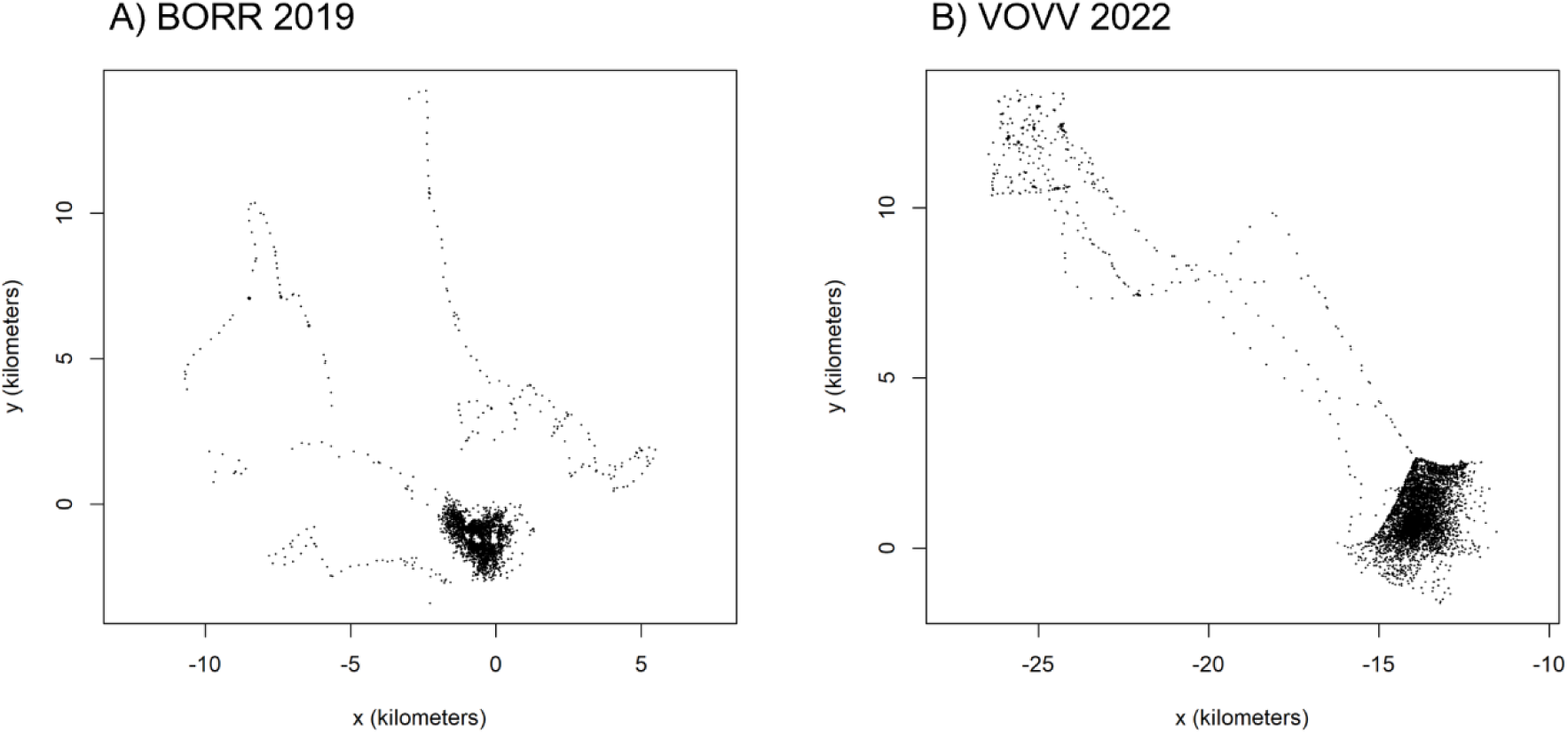
GPS locations for A) individual BORR in 2019 and B) individual VOVV in 2022, that performed a few extra-territorial trips during the study period. X and Y axis show distance from home range core.

**Figure A2.**
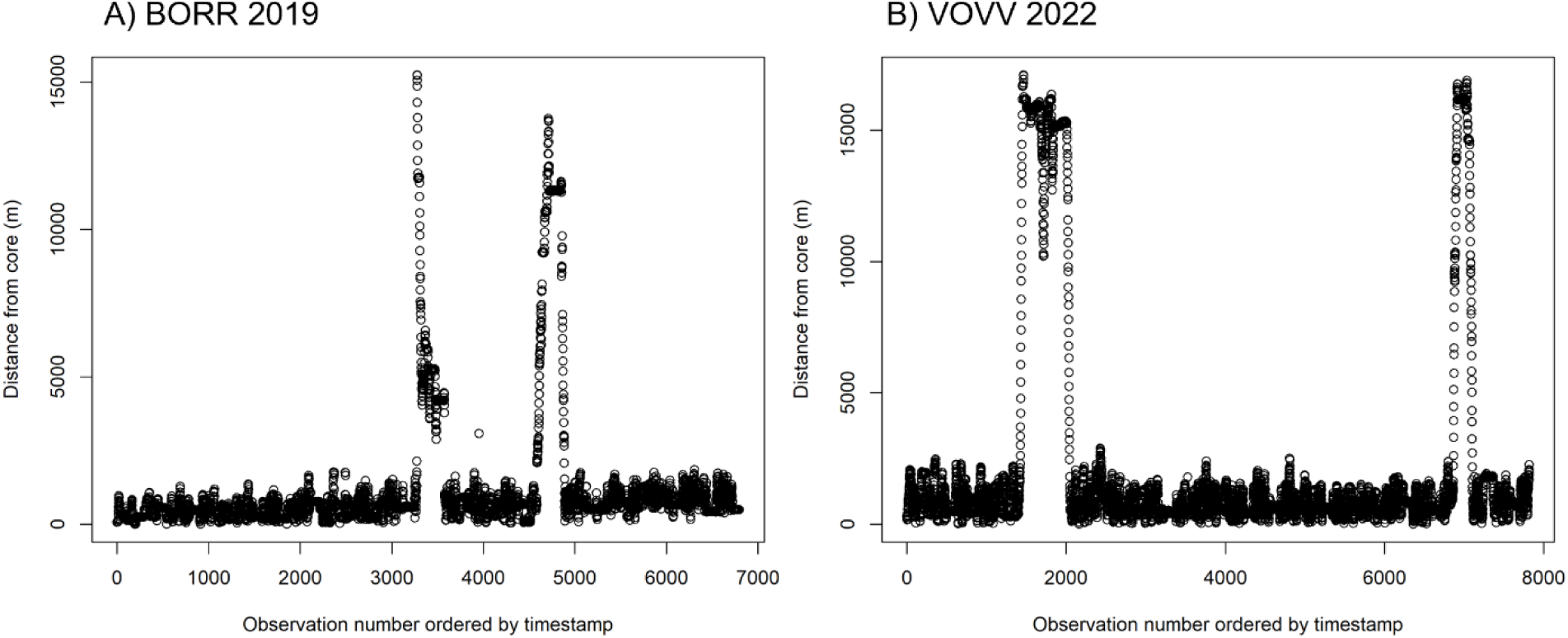
Distances between GPS locations and the core of the home range for A) individual BORR in 2019 and B) individual VOVV in 2022.

**Figure A3.**
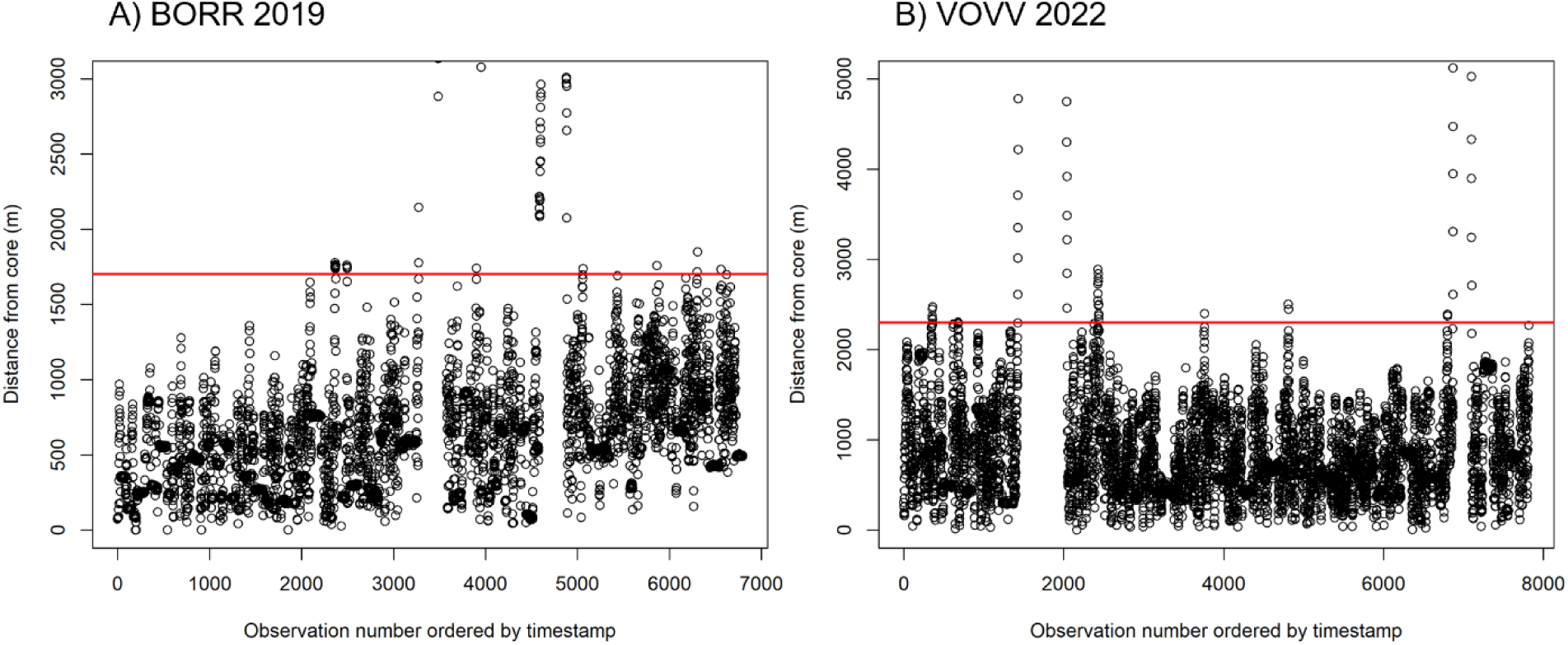
Zoom in on distances between GPS locations and the core of the home range for A) individual BORR in 2019 and B) individual VOVV in 2022. Red lines show the 1700 m and 2300 m cut-offs respectively.

**Figure A4.**
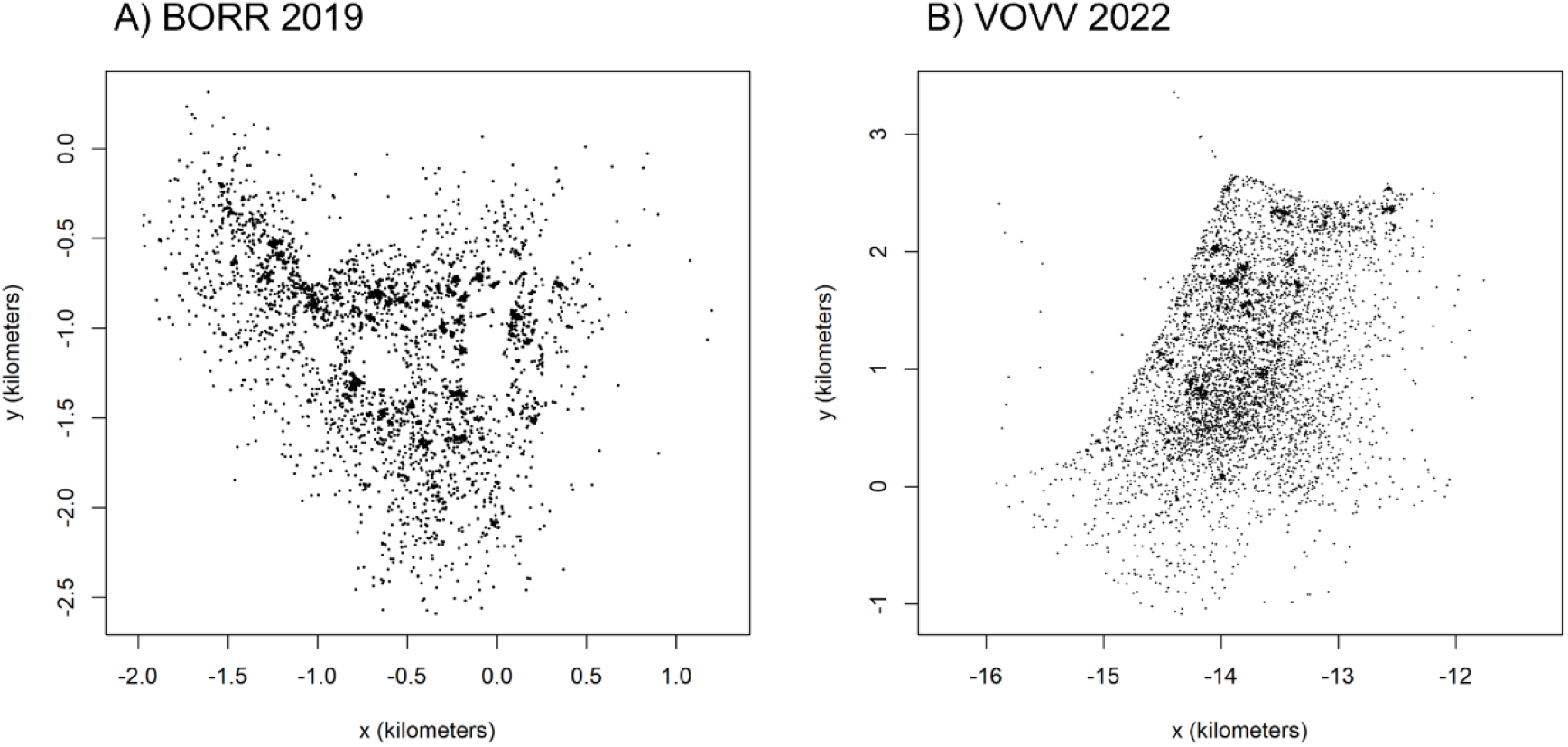
GPS locations for A) individual BORR in 2019 and 2) VOVV in 2022 after the removal of extra-territorial trips.

## Appendix B: Exclusion of datapoints near study area boundaries

Interactions with foxes that were not tracked occurred on the outer margin of our study area, thus biasing negatively our CDE estimates. For that reason, when we determined whether foxes were territorial (i.e., whether they had a part of their home range located outside of the 95% CDE), we excluded areas within 500 m of study area boundaries (Figure 2). This led to three individuals considered as non-territorial in 2019 (green female RMJJ, green male BORR and grey male BBJO, Figure 2), and one in 2022 (dark orange male BOBB, Figure 2). Using a larger buffer size of 1 km would have led to the pink female BJOV in 2022 to also be considered as non-territorial, as the 1 km buffer overlaps all her home range (Figure C1).

To test the effect of the probability of encountering a neighbour on Arctic fox behaviour, we excluded from our dataset all locations falling within the 500 m inner buffer of the study area and considered that BJOV of 2022 was territorial (results in Table 1). Due to uncertainty in BJOV territorial degree, we repeated the analyses of 2022 while excluding this individual. Results are similar to those shown in Table 1 (Table C4).

We also verified whether our choice of buffer size influenced the results by repeating models, this time excluding locations falling within an inner buffer of 1 km instead of 500 m. This also resulted in excluding all locations of individual BJOV in 2022 as they were all within the 1 km buffer (Figure C1). Results are similar to those shown in Table 1 (Table C5).

**Figure C1.**
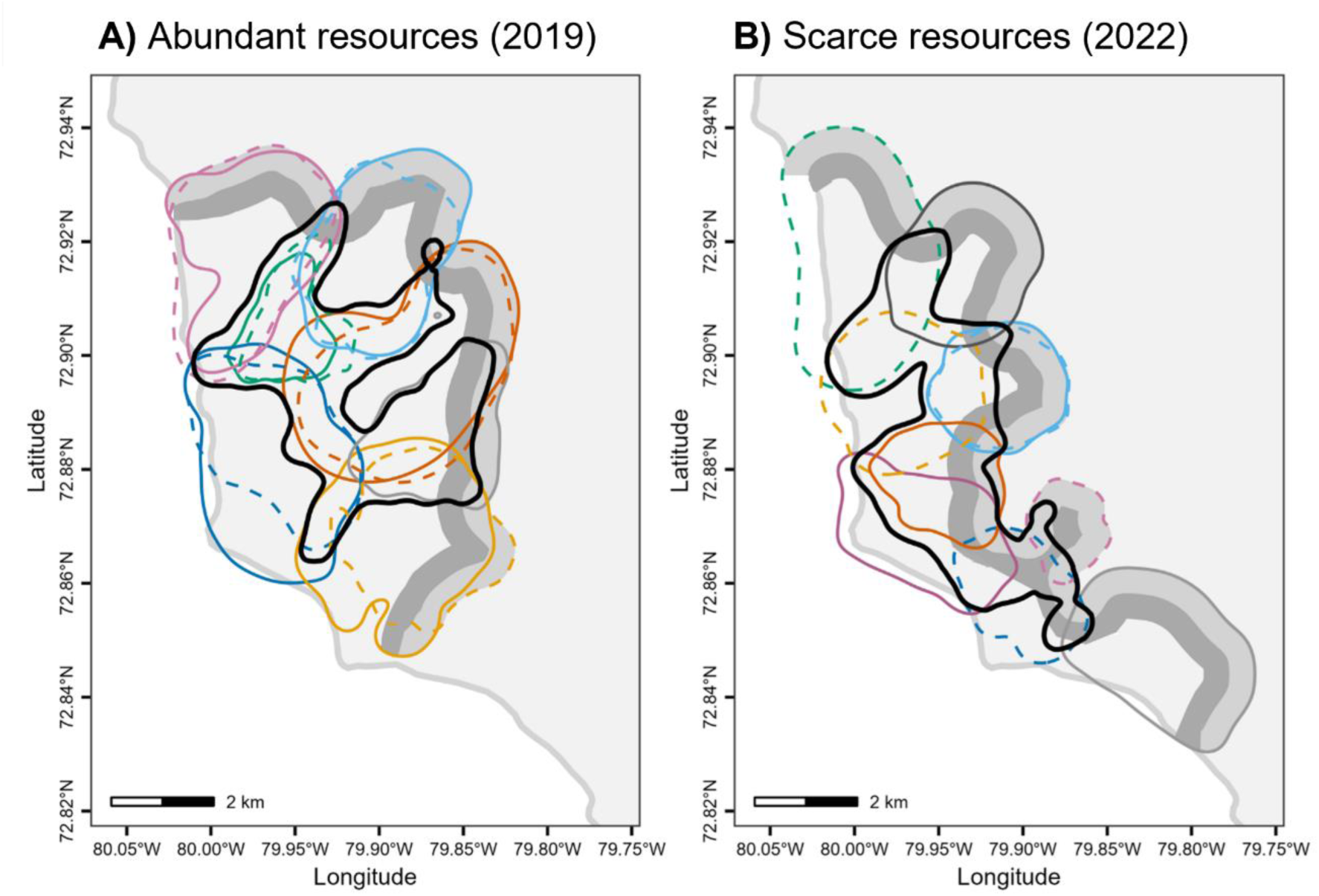
Study area with 95% auto-correlated kernel density estimation contours for A) 7 male and 6 female Arctic foxes in 2019 when resources were abundant and B) 5 male and 5 female Arctic foxes in 2022 when resources were scarce. The light grey shaded area represents a 500 m inner buffer of study area boundaries where observations were excluded, while the dark grey shaded area indicates a 1 km buffer. The thick black line is the 95% CDE contour used to determine whether foxes were territorial, and coloured lines are 95% AKDE contours. Males and females are represented by solid and dashed lines, respectively, and members of a given pair are identified by the same colour (within year).

**Table C4.** Estimates of generalized additive mixed models predicting the probability of fox digging in 2022 (n = 1500 accelerometry bursts), while excluding female BJOV for which degree of territoriality was uncertain. Model specifications are the same as those described in Table 1.

| Component | Term | Estimate | Std.Error | z.value | p.value |
| --- | --- | --- | --- | --- | --- |
| 1. Parametric coefficients | (Intercept) | -0.27 | 0.25 | -1.07 | 0.285 |
|  | Probability of encountering a neighbour | 0.070 | 0.06 | 1.14 | 0.254 |
|  | Territorial yes/no [no] | 0.292 | 0.56 | 0.52 | 0.603 |
|  | Sex [male] | -0.14 | 0.35 | -0.39 | 0.699 |
|  | Prob encounter: Territorial yes/no [no] | -0.08 | 0.17 | -0.45 | 0.655 |
| Component | Term | edf | Ref.df | Chi.sq | p.value |
| 2. Smooth terms | s(ToD) | 4.39 | 8.00 | 83.27 | <0.001 |
|  | s(day) | 1.00 | 1.00 | 3.21 | 0.073 |
|  | s(Fox ID) | 5.23 | 6.00 | 40.30 | <0.001 |
|  | s(day_factor) | 4.64 | 22.00 | 6.26 | 0.163 |

**Table C5.** Estimates of generalized additive mixed models predicting the probability of fox digging in A) 2019 (n = 1950 accelerometry bursts) and B) 2022 (n = 1500 accelerometry bursts), while excluding locations falling within a 1 km inner buffer of study area boundaries. Model specifications are the same as those described in Table 1.

| **A) Probability of fox digging when resources are abundant (2019)** |  |  |  |  |  |
| --- | --- | --- | --- | --- | --- |
| Component | Term | Estimate | Std.Error | z.value | p.value |
| 1. Parametric coefficients | (Intercept) | -0.87 | 0.34 | -2.56 | 0.011 |
|  | Probability of encountering a neighbour | -0.04 | 0.07 | -0.50 | 0.619 |
|  | Territorial yes/no [no] | 0.01 | 0.54 | 0.03 | 0.979 |
|  | Goose nest density | 0.32 | 0.06 | 5.04 | <0.001 |
|  | Sex [male] | -0.17 | 0.45 | -0.38 | 0.700 |
|  | Prob encounter: Territorial yes/no [no] | -0.47 | 0.14 | -3.29 | 0.001 |
| Component | Term | edf | Ref.df | Chi.sq | p.value |
| 2. Smooth terms | s(ToD) | 2.89 | 8.00 | 33.98 | <0.001 |
|  | s(day) | 2.77 | 3.29 | 9.20 | 0.038 |
|  | s(Fox ID) | 9.33 | 10.00 | 130.70 | <0.001 |
|  | s(day_factor) | 5.34 | 21.00 | 8.64 | 0.053 |

| **B) Probability of fox digging when resources are scarce (2022)** |  |  |  |  |  |
| --- | --- | --- | --- | --- | --- |
| Component | Term | Estimate | Std.Error | z.value | p.value |
| 1. Parametric coefficients | (Intercept) | -0.26 | 0.29 | -0.90 | 0.371 |
|  | Probability of encountering a neighbour | -0.035 | 0.07 | -0.51 | 0.608 |
|  | Territorial yes/no [no] | 0.059 | 0.65 | 0.09 | 0.927 |
|  | Sex [male] | 0.04 | 0.41 | 0.11 | 0.915 |
|  | Prob encounter: Territorial yes/no [no] | 0.07 | 0.19 | 0.38 | 0.703 |
| Component | Term | edf | Ref.df | Chi.sq | p.value |
| 2. Smooth terms | s(ToD) | 5.01 | 8.00 | 70.07 | 0.000 |
|  | s(day) | 4.54 | 5.55 | 14.89 | 0.016 |
|  | s(Fox ID) | 5.39 | 6.00 | 49.00 | 0.000 |
|  | s(day_factor) | 0.00 | 22.00 | 0.00 | 0.817 |

